# Cortex-independent open-loop control of a voluntary orofacial motor action

**DOI:** 10.1101/2021.11.15.465585

**Authors:** Michaël Elbaz, Maxime Demers, David Kleinfeld, Christian Ethier, Martin Deschênes

**Affiliations:** CERVO Research Center, Laval University, Québec City, QC G1J 2G3, Canada; Department of Physics, University of California at San Diego, La Jolla, CA 92093, USA; Section of Neurobiology, University of California at San Diego, La Jolla, CA 92093, USA

## Abstract

Whether using our eyes or our hands, we interact with our environment through mobile sensors. The efficient use of these sensory organs implies the ability to track their position; otherwise, perceptual stability and prehension would be profoundly impeded^1,2^. The nervous system may be informed about the position of a sensory organ via two complementary feedback mechanisms: peripheral reafference (external, sensory feedback) and efference copy (internal feedback)^3-6^. Yet, the potential contributions of these mechanisms remain largely unexplored. By training rats to place their vibrissae within a predetermined angular range without contact, a task that depends on knowledge of vibrissa position relative to their face, we found that peripheral reafference is not required. The presence of motor cortex is not required either, even in the absence of peripheral reafference. On the other hand, the red nucleus, which receives descending inputs from motor cortex and the cerebellum and projects to facial motoneurons^7-10^, is critical for the execution of the vibrissa task. All told, our results demonstrate the existence of an open-loop control by an internal model that is sufficient to drive voluntary motion. The internal model is independent of motor cortex and likely contains the cerebellum and associated nuclei.

## Introduction

Decoding information gathered through moving sensors – the hallmark of “active sensing” – requires keeping track of the sensors’ position^3,11^. The exploratory motor action of the vibrissae in rodents instantiates this faculty for haptic sensation. Indeed, mice and rats can report the location of an object in their vibrissa field with great precision^12-14^, which implies that they know the position of their vibrissae with respect to their face, at least during touch^15^. As facial muscles are devoid of proprioceptors^16-18^, two non-exclusive feedback mechanisms may account for knowledge of vibrissa position^3-5^: internal feedback via efference copy and sensory feedback via peripheral reafference. With efference copy, an internal copy of motor commands responsible for the movement allows the brain to keep track of the consequences of motor actions^6^. With reafferent signals, sensory receptors encode the position of the vibrissae or the kinematics of the ongoing movement^19^. Although previous studies established that both mechanisms are plausible at different anatomical levels of the vibrissa system^16,19-22^, their ethological value remains unknown.

Vibrissa tasks involving touch are not suited for disentangling the role of efference copy and reafferent signals. First, primary vibrissa afferents multiplex exafferent touch and reafferent self-motion signals^16,19,21,23^, which makes it essentially impossible to manipulate reafferent signals without interfering with the exafferent signals. Second, in the somatosensory cortex, a region that is required to localize objects with vibrissae^14,24^, there is a continuous transformation of sensory and efference copy signals along sensorimotor loops^25-28^ that blurs their respective contribution. To circumvent these limitations, we designed a vibrissa positioning task that implicitly requires knowledge of vibrissa position, or a surrogate of position such as muscle activation. Critically, the task does not involve touch and is carried out in the dark. Thus, the sensory information at play consists solely of reafferent signals that can be experimentally manipulated^5^. Altogether, our experimental model makes it possible to interrogate the existence and operating conditions of an internal model that might underpin vibrissa motor control. The internal model could implement a closed-loop comparison between executed and intended movements, as perceived by reafferent signals and efference copy, or alternatively operate in an open-loop mode, in order to adjust the motor commands.

## Results

### The vibrissa positioning task

Head-restrained rats were trained to move their left C1 vibrissa^29^ from a retracted position (70° to 90° with respect to the antero-posterior axis), denoted the “go zone”, to a protracted position (100° to 130°), denoted the “reward zone” (figure 1A). Once rats self-initiated trials by moving their vibrissa in the “go zone”, they were allowed a maximum of 10 s to reach and hold their vibrissa within the “reward zone” for a given duration. The required hold time in the reward zone adaptively increased over learning, from 10 ms initially to 1 s at the “expert” level. Once rats reached the “expert” level, the hold time was fixed and the effect of experimental manipulations was assessed. At no point in the training could the rats use touch or vision to estimate position.

**Figure 1:**
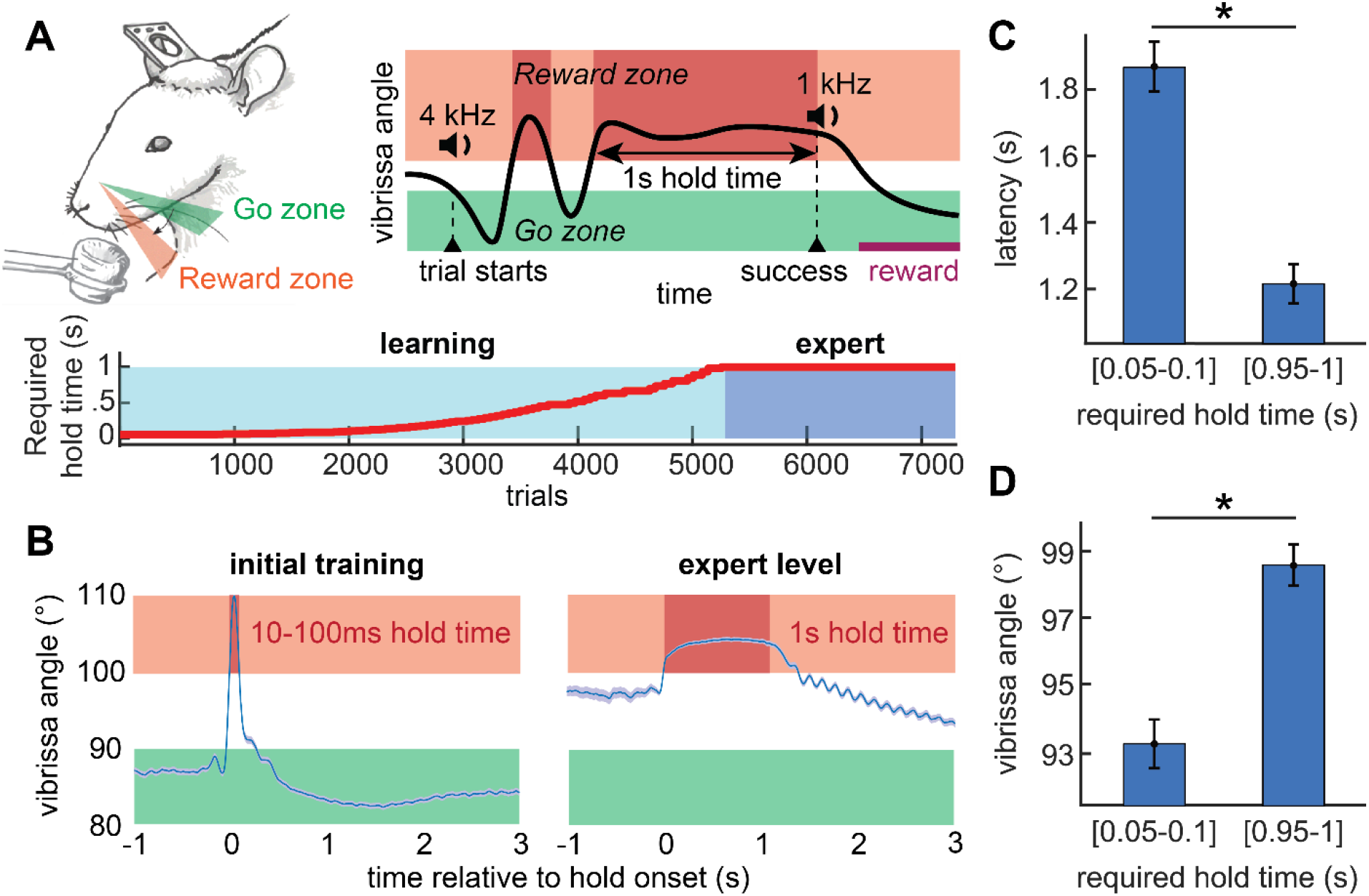
Intact animals can learn the vibrissa positioning task. A: Scheme of the vibrissa positioning task. Rats are trained to move their vibrissa from a retracted zone, i.e., the “go zone”, to a protracted zone, i.e., the “reward zone”, and maintain the vibrissa within the reward zone for a given duration, i.e., the “required hold time” (top). The required hold time adaptatively increases over training sessions (bottom; data from a representative rat). B: Mean vibrissa position over successful holds in the reward zone, over learning (mean ± 95 % confidence interval; data from a representative rat). C: Mean latency between trial onset and the time when the vibrissa is maintained in the reward zone for at least 50 ms, over learning, as the pooled mean across rats (mean ± 95 % confidence interval). D: Mean vibrissa angle over the second preceding trial onset, over learning, as the pooled mean across rats (mean ± 95 % confidence interval).

### Intact rats can learn the vibrissa positioning task

Given the short initial required hold time, naïve rats were able to obtain rewards through spontaneous vibrissa movements, such as forward twitches (figure 1B, left). Over training, rats tended to reach and maintain their vibrissa in the reward zone more swiftly following trial onset (figure 1C; mixed-effect linear regression p < 0.01). Reaching the “expert” criterion required 3760 ± 1236 trials (2.5 ± 0.8 weeks of training with 10 intact rats, mean ± 95 % CI). “Expert” animals exerted sustained protractions without rhythmic whisking of the vibrissa^30^ in the vast majority (93 % ± 1 %) of their 1-s successful hold times (figure 1B, right). As reward expectation is normally associated with whisking^31,32^, the task requires rats to curb their natural proclivity to whisk. An increasing trend to protract was also apparent during inter-trial periods (figure 1D); as a result, “expert” animals started most trials via backwards twitches (supplementary figure 1A). This suggests either that permanent sustained protraction is an effective strategy to succeed as learning progresses, or that rats did not distinguish between trials and inter-trials. We invalidate the latter hypothesis by showing that, in “expert” animals, hold times preceding premature, i.e., “false alarm”, licks are shorter during trials than during inter-trials (supplementary figure 1B; mixed-effect linear regression p < 0.01). In summary, the strategy of “expert” rats is to maintain their vibrissa close to or within the reward zone and briefly retracting it to initiate a new trial.

### Vibrissa sensory feedback is neither required before nor after learning the task

Do rats require sensory feedback to finely move their vibrissae? This is physiologically plausible since reafferent signals, encoded by mechanoreceptor afferents^19,21,33^, are present in the rodent brainstem^16,33,34^, thalamus^16,23,35^, neocortex^5,36,37^, and cerebellum^22,38^. To test the requirement of sensory feedback, we deafferented two “expert” rats, taking advantage of the separation of the vibrissa sensory and motor nerves (figure 2A and supplementary figure 2A). The deafferented rats regained their pre-lesion success rate within a few experimental sessions (7-8 sessions; figure 2B). Deafferentation did not change the vibrissa mean angle during trials (figure 2C; mixed-effect linear regression p = 0.29) nor change the across-trial variability during the “expert” holds (figures 2D and 2E; permutation test p = 0.67). Finally, the occurrence of 10°-wide holds in the reward zone was unaffected (figure 2F; mixed-effect logistic regression p = 0.42). The dispensability of a brain region in the execution of a behavior, once learnt, does not exclude its requirement for learning^39^. To test the potential requirement of sensory feedback to learn the task, we deafferented three rats before the first training session. These animals were able to learn the task and reached the “expert” level (supplementary figures 2B-E). All told, these results indicate that vibrissa sensory feedback is not required for learning nor executing the task. This implies that rats can finely control their vibrissa’s position via an open-loop controller.

**Figure 2:**
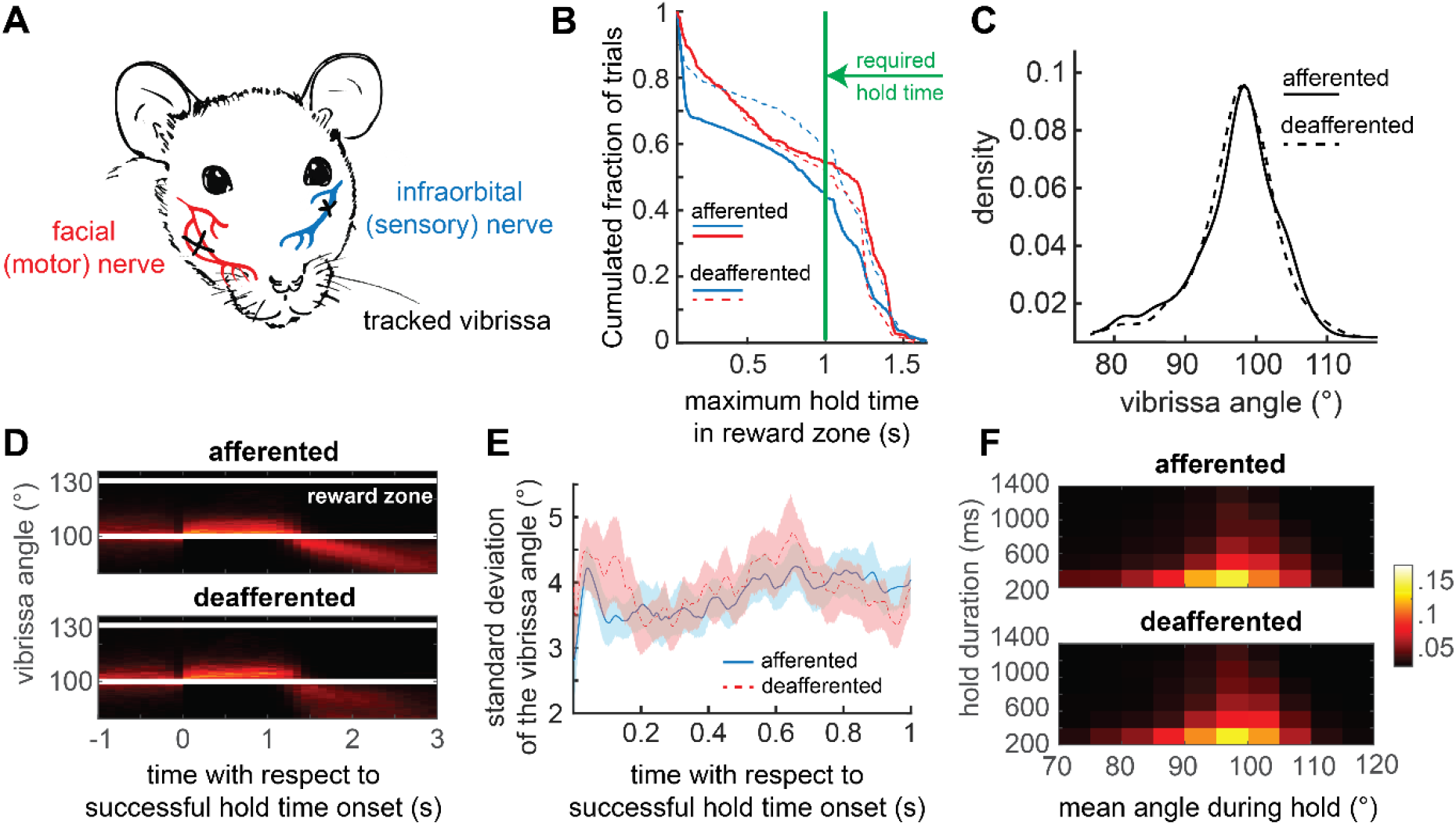
Sensory feedback is neither required before nor after learning. A: Scheme of the deafferentation procedure. The infraorbital nerve is transected on the side of the tracked vibrissa, the buccal and marginal branches of the facial nerve are transected on the opposite side. B: Empirical cumulative distribution of the maximum hold times in the reward zone across trials, before and after deafferentation. Each color represents a specific rat; required hold time: 1 s. C: Vibrissa angle distribution over trials before and after deafferentation as the pooled mean across rats; required hold time: 1 s. D: Traces of the vibrissa position during 1-s successful holds, before and after deafferentation, as the pooled mean across rats. E: Angular standard deviation across 1-s successful holds, before and after deafferentation as the pooled standard deviation across rats. F: Occurrence of 10°-wide holds before and after deafferentation as the pooled mean across “expert” rats; required hold time: 1 s. Colors represent the likelihood of occurrence.

### Motor cortex is not required to learn the task

Since vibrissa position can be decoded from the vibrissa motor cortex^20,40-42^ both in the presence and in the absence of sensory feedback^20^, we tested the necessity of motor cortex in the task, before and after deafferentation. We bilaterally ablated the motor cortex of two naïve rats (figure 3A and supplementary figure 3A) and initiated training to the task. These rats were able to learn the task and reached the “expert” level (figure 3B and supplementary figures 3B-D). We conclude that in the presence of sensory feedback, the motor cortex is not required for learning or for executing fine vibrissa movements.

**Figure 3:**
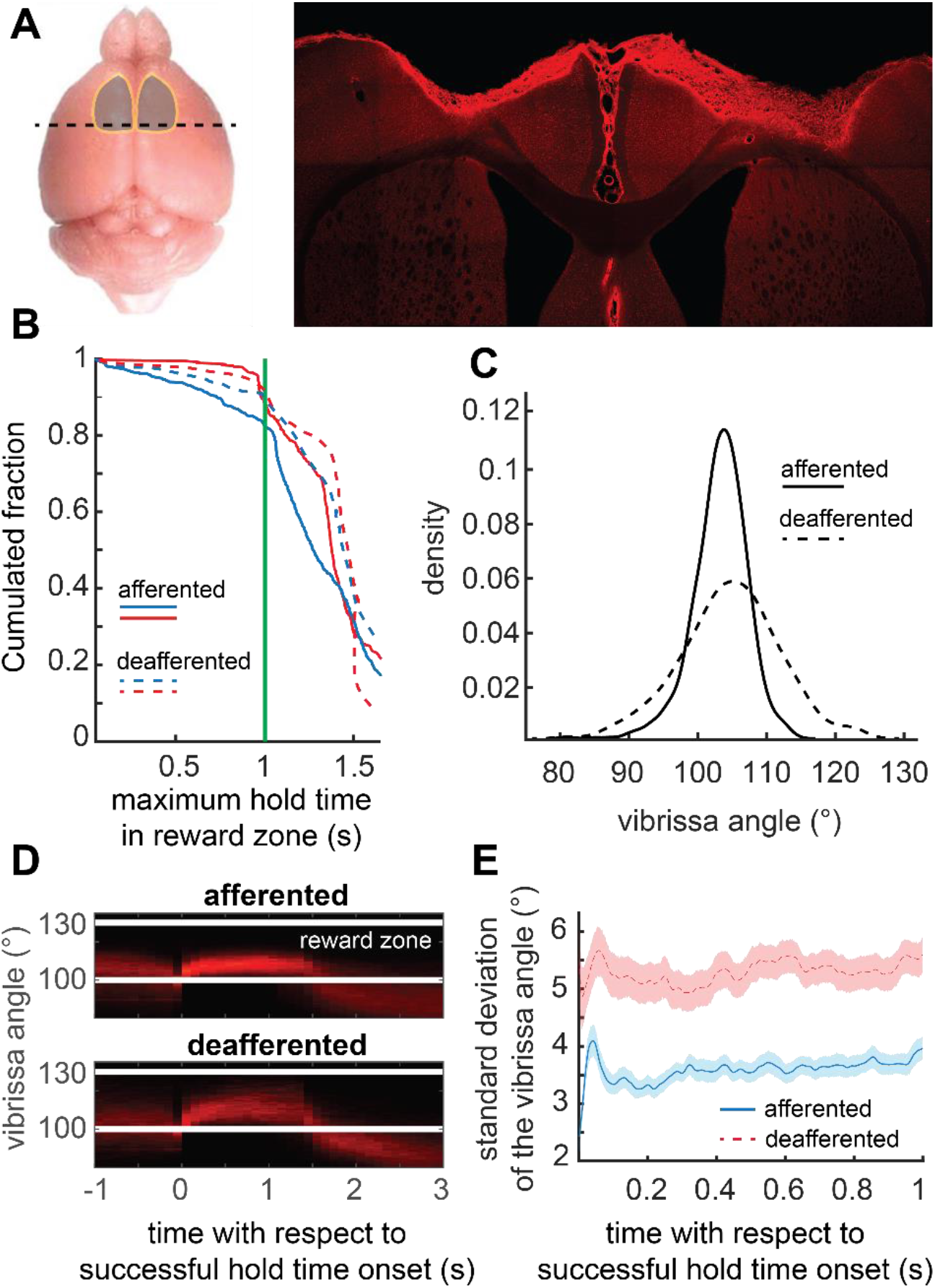
Motor cortex is not required to learn the task. A: Scheme and fluorescent microscopy image of a bilateral motor cortical lesion. The tissue was counterstained with a generic neuronal biomarker (anti-NeuN antibodies). B: Empirical cumulative distribution of the maximum hold times in the reward zone across trials in bilaterally decorticated rats, before and after deafferentation. Each color represents a specific rat; required hold time: 1 s. C: Vibrissa angle distribution over trials in bilaterally decorticated rats, before and after deafferentation as the pooled mean across rats; required hold time: 1 s. D: Traces of the vibrissa position during 1-s successful holds in bilaterally decorticated rats, before and after deafferentation as the pooled mean across rats. E: Angular standard deviation across 1-s successful holds in bilaterally decorticated rats, before and after deafferentation as the pooled standard deviation across rats.

### Absent motor cortex, sensory feedback is required for vibrissa stability

To test the requirement of motor cortex in the absence of sensory feedback, we deafferented two bilaterally decorticated “expert” rats. Both rats recovered their pre-deafferentation level in terms of occurrence of holds in the reward zone and success rate (figure 3B). Surprisingly, deafferentation of decorticated rats led to changes that did not occur in rats with intact cortex. First, the variance of the vibrissa increased during trials (figure 3C; permutation test p < 0.01) while the mean angle was unchanged (mixed-effect linear regression, p = 0.51). Second, the trial-to-trial variability in angle over the “expert” holds increased (figures 3D-E; permutation test p < 0.01). These results indicate that, in the absence of motor cortex, sensory feedback plays a non-compensable role in the ability to suppress jitter of the vibrissa within a trial and across trials. Nevertheless, in the absence of motor cortex and sensory feedback, the overall ability to voluntarily protract the vibrissa is preserved. This implies the existence of an internal model that does not require the vibrissa motor cortex or reafferent signals, but rather uses either when available to increase motor stability.

### Inactivation of the rubro-facial pathway disrupts performance

Which brain pathway might sustain the ability to perform the hold task without requiring motor cortex, but involving motor cortex at least for the case of deafferentation? Amid the numerous premotor nuclei controlling vibrissa motoneurons^7,10,43,44^, the parvocellular part of the red nucleus is of particular interest. First, it receives both cortical and cerebellar inputs^7-9^. Second, it has access to vibrissa sensory information via direct projections from the trigeminal sensory nuclei^45,46^. Third, the activity of Purkinje cells anticipates vibrissa position, even when the motor cortex is inactivated^22^. Thus, inputs to the red nucleus make it a potential integrator of reafferent and efferent signals. We examined the involvement of the rubrofacial pathway by expressing an inhibitory DREADD^47^ in rubral neurons that project to the facial motor nucleus, which drives movement of the vibrissae (figure 4A and supplementary figure 4A). This expression allowed for the conditional inactivation of rubrofacial neurons throughout the entire duration of test sessions, following intraperitoneal injection of clozapine N-oxide (CNO). To account for potential endogenous effects of CNO^48-50^, the effects of CNO were compared between rats expressing DREADD (3 rats) and rats not expressing DREADD (3 rats). Inactivation of the red nucleus dramatically impaired rats’ ability to maintain their vibrissa in the reward zone (figure 4B and supplementary figure 4B; mixed-effect logistic regression on success rates, interaction effect group:CNO p < 0.01). The vibrissa position was more retracted during trials (figure 4C; mixed-effect linear regression, interaction effect group:CNO p < 0.01), including during whisking (mixed-effect linear regression on whisking set-point, interaction effect group:CNO p < 0.05). Finally, the duration of holds within the reward zone prior to “false alarm” licks were shorter (mixed-effect linear regression, interaction effect group:CNO p < 0.01), suggesting an impeded motor perception. These results indicate that the rubrofacial pathway is critically involved in the execution of the positioning task. They imply the involvement of the red nucleus as the controller or at least relay of the internal model.

**Figure 4:**
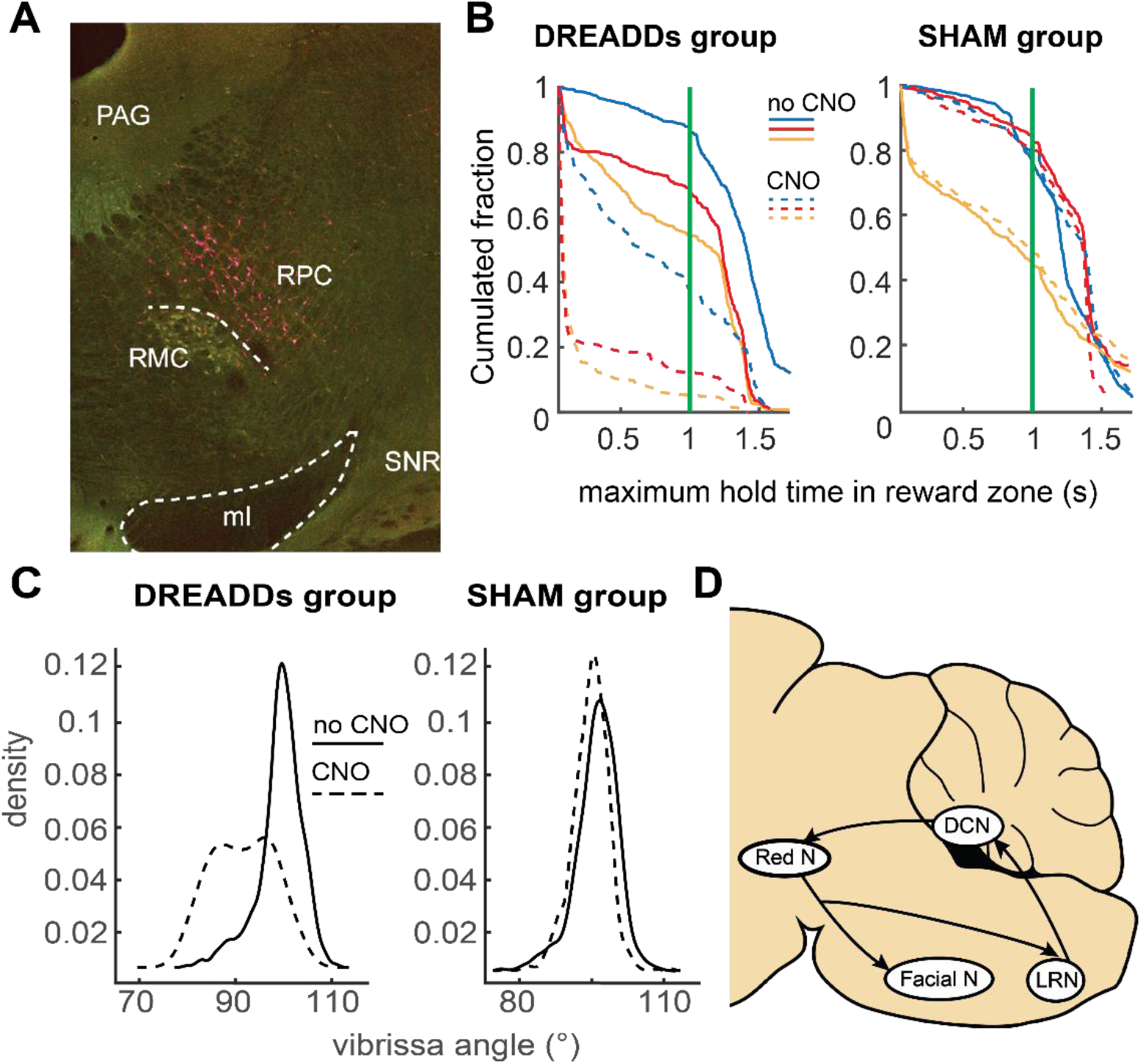
Inactivation of the rubro-facial pathway disrupts performance. A: Fluorescent microscopy image of the parvocellular red nucleus, whose neurons express an inhibitory DREADD and a red fluorescent protein, and its surroundings (PAG: periaqueductal gray; RPC: red parvocellular nucleus; RMC: red magnocellular nucleus; ML: medial lemniscus; SNR: substantia nigra pars reticulata). B: Empirical cumulative distribution of the maximum hold times in the reward zone across trials in DREADD and SHAM groups, before and after CNO administration. Each color represents a specific rat; required hold time: 1 s. C: Vibrissa angle distribution over trials in DREADD and SHAM groups, before and after CNO administration as the pooled mean across rats; required hold time: 1 s. D: Circuit diagram of rubrofacial neurons’ inputs and outputs.

## Discussion

We aimed to identify the mechanisms whereby rats can keep track of the position of their vibrissae^11,15^. Toward this goal, we trained rats to perform a vibrissa motor task without the possibility of contact (figure 1), so that deafferenting animals was strictly akin to abolishing sensory feedback. Three lessons emerged from our findings. First, rats can learn to reliably and accurately control the position of their vibrissa independently of touch and, critically, after deafferentation (figure 2). These results imply the existence of an open-loop controller and a stable motor plant. Second, rats whose motor cortex was bilaterally ablated learned the task (figure 3). Deafferentation increased motor variability in the task in decorticated but not in normal rats (figure 3). Sensory feedback and motor cortical outputs appear to compensate one another for motor stability. This implies the existence of an internal model of vibrissa position within the open-loop controller. Finally, inactivation of rubrofacial neurons drastically impeded rats’ performance (figure 4). This suggests that the red nucleus is the locus, or at least a relay station, for the internal model controller.

We transiently inactivated rather than lesioned the red nucleus, thereby testing its involvement rather than its requirement. Transient manipulations make it difficult to disambiguate the nature of the involvement, either instructive or permissive^51^, and can even lead to fast compensatory mechanisms unfolding in the space of single behavioral sessions^52^, blurring the probed normal physiology. Yet, the deficits observed in our inactivation experiments are unlikely to be strictly due to a permissive involvement of the red nucleus, since the rubrofacial neurons that we inactivated are glutamatergic premotoneurons (personal communications from Fan Wang and Jun Takato) targeting the facial nucleus, which houses only motoneurons^16,53,54^ whose single spikes trigger movements^55^. The relative retraction observed under red nucleus inactivation during both whisking and sustained movements suggest the existence of a regulatory mechanism for set-point independent of whisking. This idea is consistent with the persistence of the set-point subsequent to lesion of the whisking oscillator^56^ and, in our task, with the positive correlation between the whisking set-point and the mean angle of sustained movements adjacent to whisking fragments (supplementary figure 5).

The involvement of the red nucleus also implies the involvement of the motor cortex and/or the cerebellum, its two major inputs. Both brain regions anticipate vibrissa position^20,22^, indicating that they are part of a common network. Since cerebellar activity anticipates vibrissa position even when the motor cortex is inactivated^22^, it is tempting to speculate that the cerebellum takes control of the red nucleus in the absence of the motor cortex, possibly resulting in cerebello-rubral synaptic sprouting. The reciprocal phenomenon, cortico-rubral sprouting, has been observed at the level of the rubrospinal pathway following cerebellar lesion^57,58^. Interestingly, viral labelling reveals that rubrofacial neurons send collaterals to the lateral reticular nucleus (supplementary figure 4A and figure 4D), thus providing an anatomical substrate for an efference copy signal from the red nucleus to the cerebellum^59-61^. This might grant the cerebello-rubro-lateral reticular nucleus loop with the capability not only to control but also to correct vibrissa movements, even in the absence of sensory feedback. Several studies indicate that the cerebello-rubro-facial pathway is involved in plastic changes associated with motor learning, especially in the eye blink reflex elicited by conditional stimuli^8,62-67^. The reward-related signals to which the cerebellum has access^68-71^ likely substantiate these learning abilities.

In summary, our work establishes an experimental model to study a learned movement that is performed entirely under the control of an internal model, requiring neither sensory feedback nor motor cortex. We suggest that our work is likely to have a significant impact in facilitating the study of internal models in mammals.

## Methods

This study’s protocol was approved by the Comité de Protection des Animaux de l’Université Laval (CPAUL). All procedures were carried out in strict accordance with the Canadian animal care and use guidelines. All surgical procedures were performed under ketamine-xylazine anesthesia.

### Animals

Eighteen male Long Evans rats, 250 to 350 g in mass (Charles River), were used for combined behavioral, neurophysiological and anatomical experiments. They were housed in a reverse dark-light cycle in a facility with controlled temperature and humidity. After a week of daily handling aimed at getting them habituated to the experimental room and to the experimenter, they were implanted with a plate for head fixation, following procedures previously described^72^. A week after head-plate implantation, the rats were placed under water restriction. They were head-restrained over increased periods of time and given water concomitantly. After being habituated and comfortable enough to drink enough water while being head-restrained (10 ml/100 g body weight/day), we began exposing them to the vibrissa positioning task (referred to below as “the task”). From that moment on, all their vibrissae but left C1 were trimmed weekly under light isoflurane anesthesia in order to optimize the online detection of the vibrissa of interest and to prevent tactile contact with any element of the surrounding environment. The choice of C1 was motivated by the relative stable dorso-ventral (azimuthal) level of this vibrissa along its whole retraction-protraction range. Rats were trained to the task twice a day (at least 4 hours apart between both sessions), 20 minutes per session, from Monday to Friday.

### Vibrissa positioning task

All experiments were carried out with head-restrained rats in a silent and dark room under infrared illumination (880 nm). The task was implemented with custom MATLAB (Mathworks) scripts operating two cameras: one used as a lickometer to detect tongue movements, the other one to detect vibrissa position with respect to the head antero-posterior axis (i.e., monitoring the absolute vibrissa angle).

Task trials were self-initiated by the rat when its vibrissa was detected in the “go zone” (70 to 90° with respect to the head axis). Trial onset was accompanied by an auditory cue (4 kHz, 300 ms), after which the rat had 10 seconds to position its vibrissa in the “reward zone” (100 to 130°) and maintain it for a given required hold time. Over training, the required hold time increased from 10 ms up to 1 s in an adaptive fashion, depending on the success rate of previously “attempted trials”. An attempted trial is defined as a trial during which the vibrissa goes 5° beyond the upper limit of the go zone, i.e., beyond 95°. When the mean success rate of the fifty past attempted trials reached 50%, the hold time was increased by 5 % of its previous value, up to 1 s (expert level). At the end of each trial, a 1 kHz or 8 kHz auditory signal (duration: 300 ms) respectively indicated whether the trial was successful or not. In case of success, a pump was activated to deliver 75 µl of water. After a successful trial, a pause of variable duration (5 - 8 s), during which it was impossible to initiate another trial, allowed animals to lick the delivered water without interfering with the task. After an unsuccessful trial, there was a 1-s pause to prevent the juxtaposition of trials in case the vibrissa was in the go zone when a trial ended, which allowed us to play distinctly the sounds announcing the end of the failed trial and the onset of the following one.

### Deafferentation

On the side of the tracked vibrissa (left), we transected the infraorbital (sensory) branch of the infraorbital nerve at its entrance in the orbit; on the opposite side (right), we transected the branches of the facial motor nerve innervating the musculature of the mystacial pad (namely, the buccal and marginal branches). Thus, the mystacial pad did not convey any information related to self-motion. Four days after deafferentation, rats were re-exposed to the task. The effectiveness of the facial nerve lesion was assessed by the absence of vibrissa movement on the corresponding side. At the end of behavioral experiments, the effectiveness of the deafferentation was assessed by the absence of an evoked local field potential in the ventral posteromedial nucleus of the thalamus upon electrical stimulation of the mystacial pad (supplementary figure 2A). The recording site was labeled by an iontophoretic injection of Chicago Sky Blue (Sigma-Aldrich). Thereafter the rat was perfused, and brain tissue was processed for histology.

### Cortical lesion

The vibrissa motor cortex was unilaterally or bilaterally lesioned by the application, over the pia mater, of a small crystal of silver nitrate, a strong cauterizing agent^73^ (centered 2 mm on the antero-posterior axis and 2 mm on the medio-lateral axis with respect to the bregma). The crystal was left in place for 5 min to allow diffusion of the chemical to the deep layers. Then, the cortex was abundantly rinsed with saline and sucked out. At the end of behavioral experiments, the rat was perfused and brain tissue was cut at 60 microns on a freezing microtome. Sections were immunoreacted with a rabbit anti-NeuN antibody (Invitrogen) and an anti-rabbit antibody conjugated to Alexa 594 (Thermofisher). Images of the cortical lesion were acquired using a slide scanner (Huron Digital Pathology).

### Transient inactivation of the red nucleus

Inhibitory DREADD was expressed in rubrofacial neurons via dual viral injections (100 nanoliters each); AAV-hSyn-DIO-hM4D(Gi)-mCherry (Addgene #44362) was injected in the right parvocellular red nucleus and retrogradeAAV-hSyn.Cre.WPRE.hGH (Addgene #105553) was injected in the lateral sector of the left facial nucleus. The red nucleus was located by stereotaxy (1 mm lateral to the midline, 5.5 mm behind the bregma, 6.5 mm below the dura). To target injections in the facial nucleus we first used micro-stimulation to elicit vibrissa deflection. Thereafter the virus was injected at the very same location. After the injections, a head-plate was fixed to the skull with screws and acrylic cement. Inactivation experiments were carried out at least 4 weeks after the viral injections. To inactivate rubrofacial neurons expressing DREADD, Clozapine N-oxide dihydrochloride (Tocris) was intraperitoneally injected (2 mg/kg) one hour prior to the behavioral test. The exact same procedure was employed for rats who did not express DREADD (sham group). When all the behavioral sessions were completed, the animals were perfused and brains were processed for histology. Images of the red nucleus and brainstem containing labelled neurons were acquired using a confocal microscope (Zeiss).

### Data analysis

All data analyses were carried out under MATLAB (Mathworks). All analyses on groups of animals were pooled so that individual animals’ contribution to the final measure was equivalent. All error bars and shaded plot areas correspond to 95 % confidence intervals.

#### Licking learning

To evaluate learning of the vibrissa positioning task, vibrissa data was analyzed only after rats had learnt to lick reliably upon water delivery, i.e., after the reward could be tightly associated with a preceding action. The trial from which a rat is considered to lick reliably is defined as the first trial preceded by at least 80% licking occurrence in the first 2 s of the latest 250 post-reward pauses. Reliable licking upon reward delivery was achieved after 603 ± 191 rewards (mean ± 95 % CI).

#### Task learning

As per our behavioral protocol, trial completion and reward delivery occurred as soon as rats maintained their vibrissa in the reward zone for the required hold time. As required hold time increased over learning, a direct comparison of the vibrissa position over entire trials at different stages of learning is not possible. To compute the latency between trial onset and holds in the reward zone of a duration greater than or equal to 50 ms, we excluded trials whose required hold time was lower than 50 ms. In contrast, periods between trials, i.e., “inter-trials”, are similar over learning: they all end as soon as the vibrissa reaches the go zone, provided that inter-trial pause has elapsed. Thus, vibrissa position during inter-trials can be compared over learning. In analyses of inter-trials, pauses following successful trials dedicated to licking were excluded from the analyses as well as the 1-s pause following unsuccessful trials.

#### “False alarm” licks at expert level

These are licks which did not follow water delivery. They occurred either during trials or during inter-trials; in this latter case, outside the dedicated licking period following successful trials. We computed the duration of the longest sustained hold within the reward zone in the 1-s window preceding every “false alarm” lick’s occurrence. To compare these hold times between trials and inter-trials, we excluded licks occurring during trials when an incursion in the go zone occurred in the preceding 1-s window because such an incursion would not have been possible during inter-trials, leading to their end and triggering a new trial.

#### Recovery from deafferentation

The session from which rats were considered to have recovered from deafferentation was the first of the three consecutive sessions whose pooled success rate was not statistically different from the pooled success rate of the last three sessions preceding deafferentation (chi-squared test, p > 0.05). All subsequent analyses aimed at comparing steady-state execution before and after deafferentation were carried out with the post-recovery data.

#### Spectral analyses

All spectral analyses were carried out using multitaper estimates^74^, as implemented in the Chronux MATLAB toolbox^75^. Whisking bouts of over 500 ms duration were detected using spectrograms, and whisking parameters (set-point, amplitude and phase) were extracted using the Hilbert transform^20,76^.

#### Whisking set-point and adjacent sustained holds

For expert rats, we regressed the angle of adjacent sustained movements as a function of the whisking set-point, including every trial containing whisking bout(s) and at least 500-ms long 10°-wide hold(s).

## Figures

**Supplementary Figure 1:**
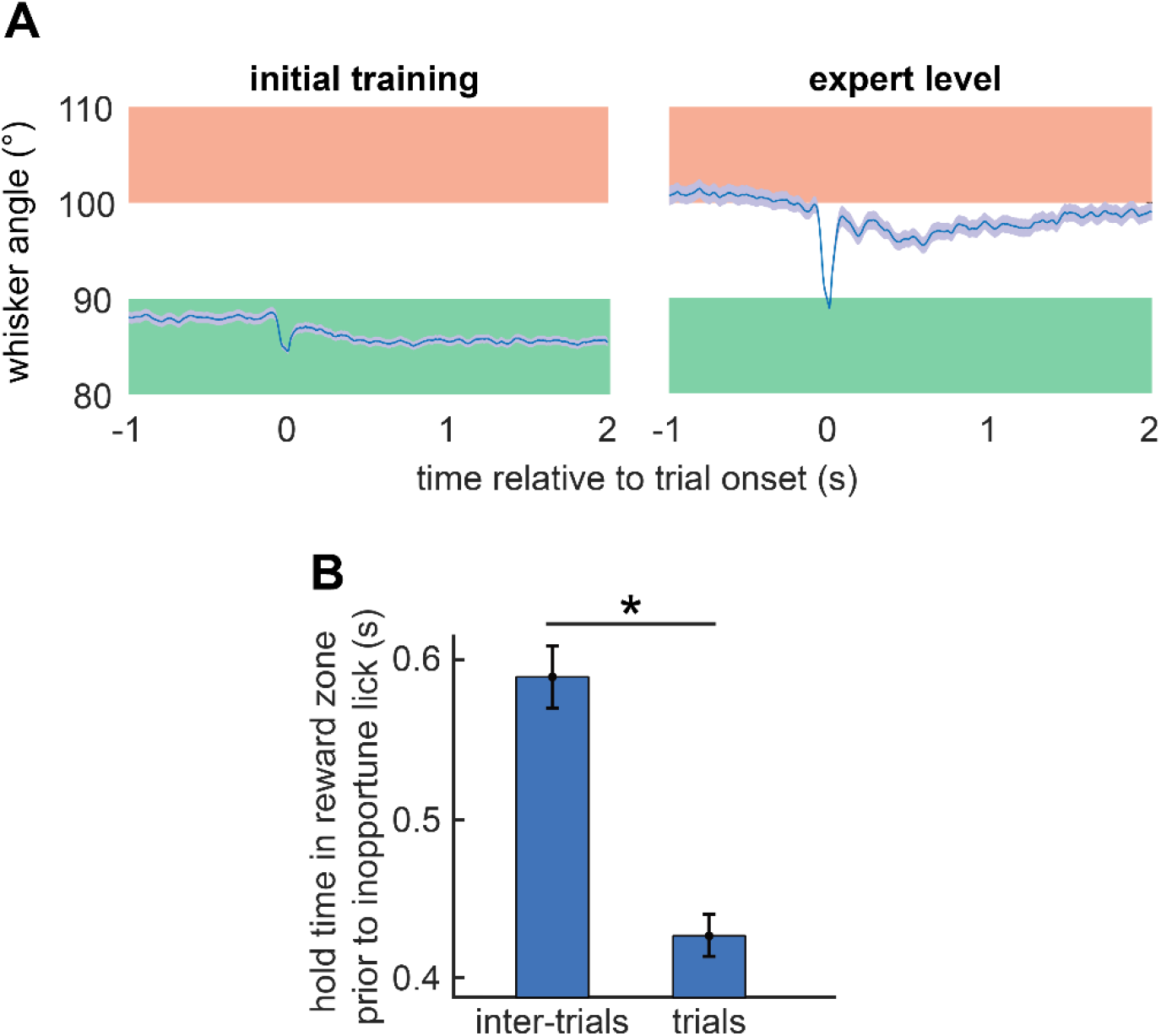
Intact animals can learn the vibrissa positioning task. A: Mean vibrissa position at trial initiation, throughout learning (mean ± 95 % confidence interval; data from a representative rat). B: Hold times in reward zone preceding “false alarm” licks during trials vs. inter-trials, as the pooled mean across rats; required hold time: 1 s.

**Supplementary Figure 2:**
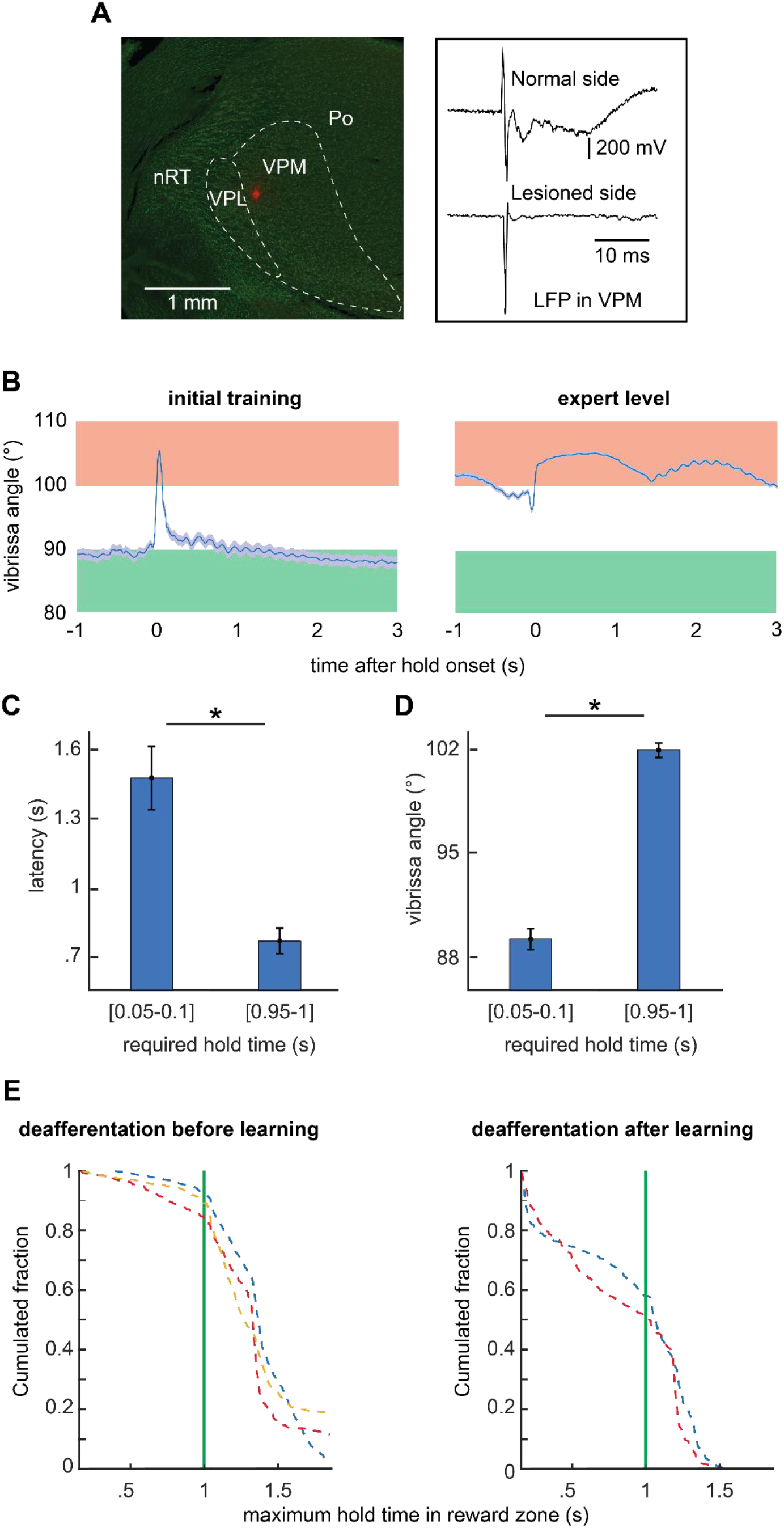
Sensory feedback is neither required before nor after learning. A: Post-hoc assessment of the effectiveness of the infraorbital nerve lesion, through bilateral long field potential (LFP) recording in the vibrissa ventral posteromedial nucleus (VPM) of the thalamus, during electrical stimulation of pad muscles (data from a representative rat). The fluorescent red spot in the VPM corresponds to a Chicago Sky Blue spot, iontophoretically injected at the end of the LFP recording contralaterally to the infraorbital lesion (VPL: ventral posterolateral nucleus of the thalamus; PO: posterior nucleus of the thalamus; nRT: reticular nucleus of the thalamus). B: Mean vibrissa position over successful holds in the reward zone throughout learning, for a deafferented rat. C: Mean latency between trial onset and the time when the vibrissa is maintained in the reward zone for at least 50 ms, over learning, as the pooled mean across deafferented rats (mean ± 95 % confidence interval). D: Mean vibrissa angles over the second preceding trial onset over learning, as the pooled mean across deafferented rats (mean ± 95 % confidence interval). E: Empirical cumulative distribution of the maximum hold times in the reward zone across trials in rats deafferented before and after learning. Each color represents a specific rat; required hold time: 1 s.

**Supplementary Figure 3:**
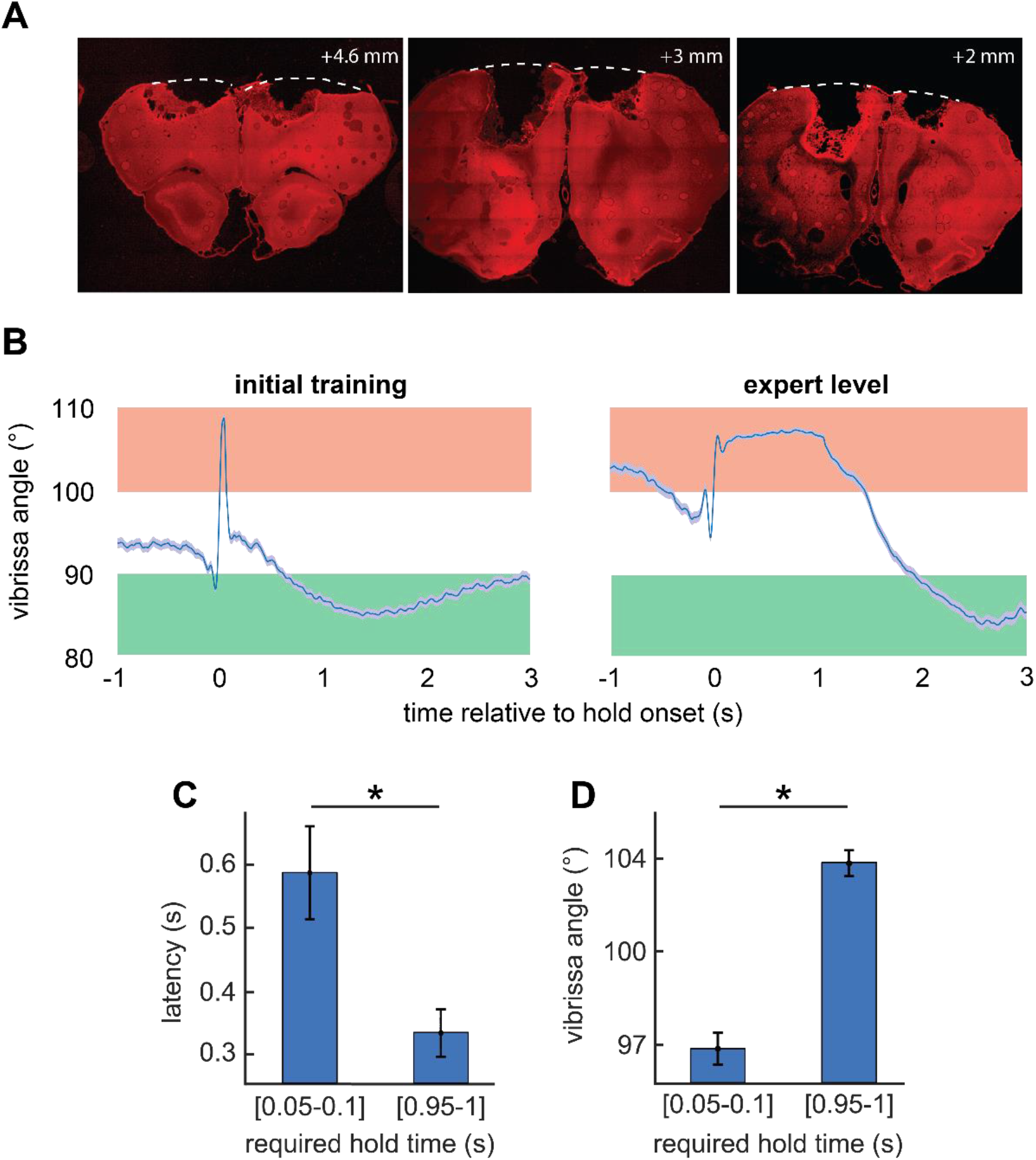
Motor cortex is not required to learn the task. A: Fluorescent microscopy images of a bilateral motor cortical lesion. Neurons were counterstained with anti-NeuN antibodies. B: Mean vibrissa position over successful holds in the reward zone throughout learning, for a bilaterally decorticated rat (mean ± 95 % confidence interval). C: Mean latency between trial onset and the time when the vibrissa is maintained in the reward zone for at least 50 ms, over learning, as the pooled mean across bilaterally decorticated rats (mean ± 95 % confidence interval). D: Mean vibrissa angles over the second preceding trial onset over learning, as the pooled mean for bilaterally decorticated rats (mean ± 95 % confidence interval).

**Supplementary Figure 4:**
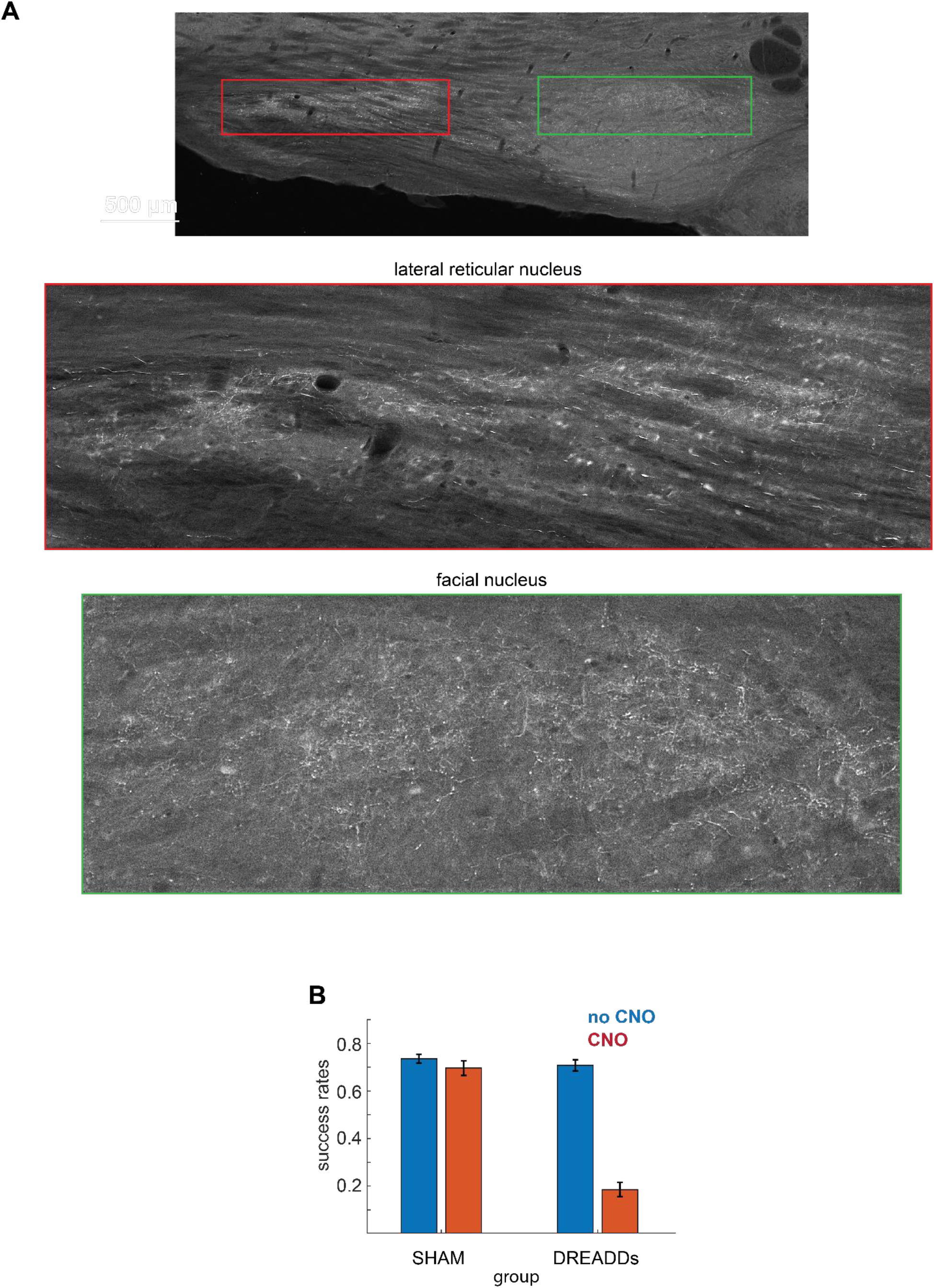
Inactivation of the rubro-facial pathway disrupts performance. A: Collaterals of rubrofacial neurons revealed by fluorescent microscopy. The labelled neurons coexpress a red fluorescent protein and an inhibitory DREADD allowing for their conditional inactivation. B: Success rates during trials in DREADD and SHAM groups, before and after CNO administration (pooled mean across rats; required hold time: 1 s).

**Supplementary Figure 5.**
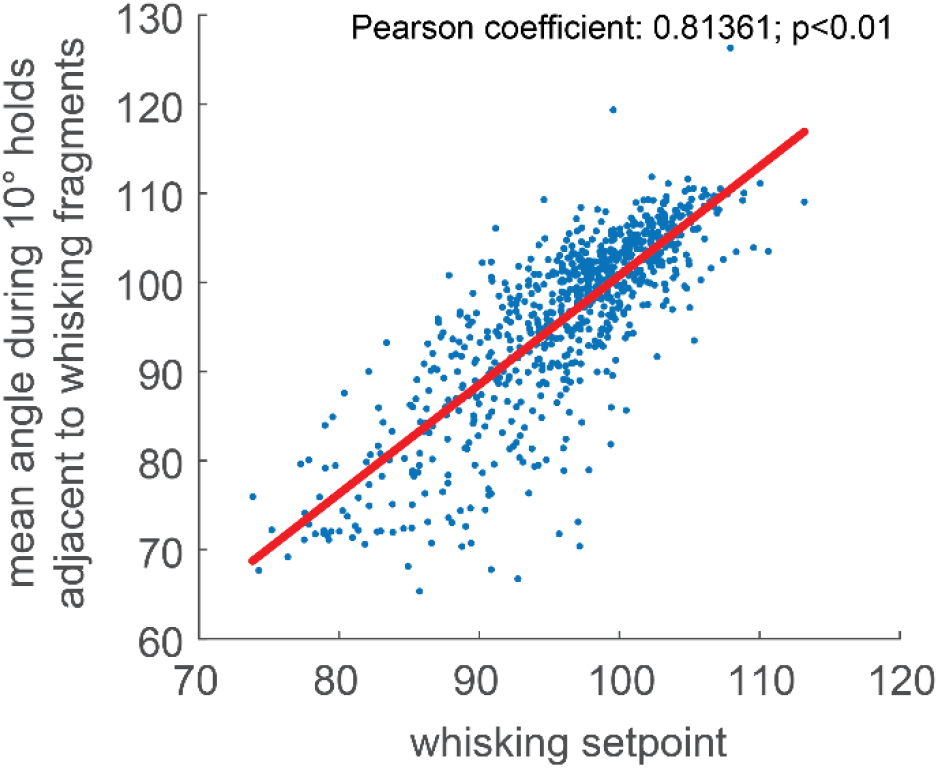
Linear relationship between the whisking setpoint and the mean angle of the vibrissa angle during adjacent 10°-wide 500-ms long holds, over trials (data from a representative rat).

## Acknowledgements

We thank Ehud Ahissar (Weizmann Institute) for discussions, Andrew Miri (Northwestern) and Windsor Ting (Laval University) for comments on the manuscript, Jun Takato (MIT) and Fan Wang (MIT) for sharing unpublished data, Sergiu Ftomov (Laval University) for technical help, and Gabrielle Lahaye for preparing illustrations.

## Author contributions

M. Deschênes and M.E. devised the project with the help of C.E. to design the behavioral task; M. Demers, M. Deschênes and M.E. carried out the surgical procedures; M.E. carried out the behavioral experiments; M.E. analyzed data with assistance from C.E. and D.K.; M.E. wrote the manuscript with assistance from M. Deschênes, C.E. and D.K.

## Additional information

Supplementary Information is available for this paper.

Correspondence and requests for materials should be addressed to Martin Deschênes or Michaël Elbaz.

Reprints and permissions information is available at www.nature.com/reprints

## References

1 Connolly, J. D. & Goodale, M. A. The role of visual feedback of hand position in the control of manual prehension. Exp Brain Res 125, 281–286, doi:10.1007/s002210050684 (1999).

2 Wurtz, R. H. Corollary Discharge Contributions to Perceptual Continuity Across Saccades. Annu Rev Vis Sci 4, 215–237, doi:10.1146/annurev-vision-102016-061207 (2018).

3 Wolpert, D. M. & Ghahramani, Z. Computational principles of movement neuroscience. Nat Neurosci 3 Suppl, 1212–1217, doi:10.1038/81497 (2000).

4 Wolpert, D. M., Ghahramani, Z. & Jordan, M. I. An internal model for sensorimotor integration. Science 269, 1880–1882, doi:10.1126/science.7569931 (1995).

5 Fee, M. S., Mitra, P. P. & Kleinfeld, D. Central versus peripheral determinants of patterned spike activity in rat vibrissa cortex during whisking. J Neurophysiol 78, 1144–1149, doi:10.1152/jn.1997.78.2.1144 (1997).

6 Crapse, T. B. & Sommer, M. A. Corollary discharge across the animal kingdom. Nat Rev Neurosci 9, 587–600, doi:10.1038/nrn2457 (2008).

7 Hattox, A. M., Priest, C. A. & Keller, A. Functional circuitry involved in the regulation of whisker movements. J Comp Neurol 442, 266–276, doi:10.1002/cne.10089 (2002).

8 Pacheco-Calderon, R., Carretero-Guillen, A., Delgado-Garcia, J. M. & Gruart, A. Red nucleus neurons actively contribute to the acquisition of classically conditioned eyelid responses in rabbits. J Neurosci 32, 12129–12143, doi:10.1523/JNEUROSCI.1782-12.2012 (2012).

9 Daniel, H., Billard, J. M., Angaut, P. & Batini, C. The interposito-rubrospinal system. Anatomical tracing of a motor control pathway in the rat. Neuroscience Research 5, 87–112, doi:10.1016/0168-0102(87)90027-7 (1987).

10 Isokawa-Akesson, M. & Komisaruk, B. R. Difference in projections to the lateral and medial facial nucleus: anatomically separate pathways for rhythmical vibrissa movement in rats. Exp Brain Res 65, 385–398, doi:10.1007/BF00236312 (1987).

11 Kleinfeld, D. & Deschenes, M. Neuronal basis for object location in the vibrissa scanning sensorimotor system. Neuron 72, 455–468, doi:10.1016/j.neuron.2011.10.009 (2011).

12 Knutsen, P. M., Pietr, M. & Ahissar, E. Haptic object localization in the vibrissal system: behavior and performance. J Neurosci 26, 8451–8464, doi:10.1523/JNEUROSCI.1516-06.2006 (2006).

13 Mehta, S. B., Whitmer, D., Figueroa, R., Williams, B. A. & Kleinfeld, D. Active spatial perception in the vibrissa scanning sensorimotor system. PLoS Biol 5, e15, doi:10.1371/journal.pbio.0050015 (2007).

14 O’Connor, D. H. et al. Vibrissa-based object localization in head-fixed mice. J Neurosci 30, 1947–1967, doi:10.1523/JNEUROSCI.3762-09.2010 (2010).

15 Cheung, J., Maire, P., Kim, J., Sy, J. & Hires, S. A. The Sensorimotor Basis of Whisker-Guided Anteroposterior Object Localization in Head-Fixed Mice. Curr Biol, doi:10.1016/j.cub.2019.07.068 (2019).

16 Moore, J. D., Mercer Lindsay, N., Deschenes, M. & Kleinfeld, D. Vibrissa Self-Motion and Touch Are Reliably Encoded along the Same Somatosensory Pathway from Brainstem through Thalamus. PLoS Biol 13, e1002253, doi:10.1371/journal.pbio.1002253 (2015).

17 Bowden, R. E. & Mahran, Z. Y. The functional significance of the pattern of innervation of the muscle quadratus labii superioris of the rabbit, cat and rat. J Anat 90, 217–227 (1956).

18 Kleinfeld, D., Berg, R. W. & O’Connor, S. M. Anatomical loops and their electrical dynamics in relation to whisking by rat. Somatosens Mot Res 16, 69–88, doi:10.1080/08990229970528 (1999).

19 Severson, K. S., Xu, D., Yang, H. & O’Connor, D. H. Coding of whisker motion across the mouse face. Elife 8, doi:10.7554/eLife.41535 (2019).

20 Hill, D. N., Curtis, J. C., Moore, J. D. & Kleinfeld, D. Primary motor cortex reports efferent control of vibrissa motion on multiple timescales. Neuron 72, 344–356, doi:10.1016/j.neuron.2011.09.020 (2011).

21 Severson, K. S. et al. Active Touch and Self-Motion Encoding by Merkel Cell-Associated Afferents. Neuron 94, 666–676 e669, doi:10.1016/j.neuron.2017.03.045 (2017).

22 Chen, S., Augustine, G. J. & Chadderton, P. The cerebellum linearly encodes whisker position during voluntary movement. Elife 5, e10509, doi:10.7554/eLife.10509 (2016).

23 Gutnisky, D. A. et al. Mechanisms underlying a thalamocortical transformation during active tactile sensation. PLoS Comput Biol 13, e1005576, doi:10.1371/journal.pcbi.1005576 (2017).

24 Guo, Z. V. et al. Flow of cortical activity underlying a tactile decision in mice. Neuron 81, 179–194, doi:10.1016/j.neuron.2013.10.020 (2014).

25 Petreanu, L. et al. Activity in motor-sensory projections reveals distributed coding in somatosensation. Nature 489, 299–303, doi:10.1038/nature11321 (2012).

26 Mao, T. et al. Long-range neuronal circuits underlying the interaction between sensory and motor cortex. Neuron 72, 111–123, doi:10.1016/j.neuron.2011.07.029 (2011).

27 Veinante, P. & Deschenes, M. Single-cell study of motor cortex projections to the barrel field in rats. J Comp Neurol 464, 98–103, doi:10.1002/cne.10769 (2003).

28 Ahissar, E. & Kleinfeld, D. Closed-loop neuronal computations: focus on vibrissa somatosensation in rat. Cereb Cortex 13, 53–62, doi:10.1093/cercor/13.1.53 (2003).

29 Brecht, M., Preilowski, B. & Merzenich, M. M. Functional architecture of the mystacial vibrissae. Behavioural Brain Research 84, 81–97, doi:10.1016/s0166-4328(97)83328-1 (1997).

30 Welker. Analysis of Sniffing of the Albino Rat. (1964).

31 Dominiak, S. E. et al. Whisking Asymmetry Signals Motor Preparation and the Behavioral State of Mice. J Neurosci 39, 9818–9830, doi:10.1523/JNEUROSCI.1809-19.2019 (2019).

32 Clarke, S. & Trowill, J. A. Sniffing and motivated behavior in the rat. Physiol Behav 6, 49–52, doi:10.1016/0031-9384(71)90013-8 (1971).

33 Wallach, A., Bagdasarian, K. & Ahissar, E. On-going computation of whisking phase by mechanoreceptors. Nat Neurosci 19, 487–493, doi:10.1038/nn.4221 (2016).

34 Zucker, E. & Welker, W. I. Coding of somatic sensory input by vibrissae neurons in the rat’s trigeminal ganglion. Brain Res 12, 138–156, doi:10.1016/0006-8993(69)90061-4 (1969).

35 Urbain, N. et al. Whisking-Related Changes in Neuronal Firing and Membrane Potential Dynamics in the Somatosensory Thalamus of Awake Mice. Cell Rep 13, 647–656, doi:10.1016/j.celrep.2015.09.029 (2015).

36 Cheung, J. A. et al. Independent representations of self-motion and object location in barrel cortex output. PLoS Biol 18, e3000882, doi:10.1371/journal.pbio.3000882 (2020).

37 Berg, R. W. & Kleinfeld, D. Rhythmic whisking by rat: retraction as well as protraction of the vibrissae is under active muscular control. J Neurophysiol 89, 104–117, doi:10.1152/jn.00600.2002 (2003).

38 O’Connor, S. M., Berg, R. W. & Kleinfeld, D. Coherent electrical activity between vibrissa sensory areas of cerebellum and neocortex is enhanced during free whisking. J Neurophysiol 87, 2137–2148, doi:10.1152/jn.00229.2001 (2002).

39 Kawai, R. et al. Motor cortex is required for learning but not for executing a motor skill. Neuron 86, 800–812, doi:10.1016/j.neuron.2015.03.024 (2015).

40 Ebbesen, C. L., Doron, G., Lenschow, C. & Brecht, M. Vibrissa motor cortex activity suppresses contralateral whisking behavior. Nat Neurosci 20, 82–89, doi:10.1038/nn.4437 (2017).

41 Sreenivasan, V. et al. Movement Initiation Signals in Mouse Whisker Motor Cortex. Neuron 92, 1368–1382, doi:10.1016/j.neuron.2016.12.001 (2016).

42 Friedman, W. A., Zeigler, H. P. & Keller, A. Vibrissae motor cortex unit activity during whisking. J Neurophysiol 107, 551–563, doi:10.1152/jn.01132.2010 (2012).

43 Takatoh, J. et al. New modules are added to vibrissal premotor circuitry with the emergence of exploratory whisking. Neuron 77, 346–360, doi:10.1016/j.neuron.2012.11.010 (2013).

44 Sreenivasan, V., Karmakar, K., Rijli, F. M. & Petersen, C. C. Parallel pathways from motor and somatosensory cortex for controlling whisker movements in mice. Eur J Neurosci 41, 354–367, doi:10.1111/ejn.12800 (2015).

45 Godefroy, J. N., Thiesson, D., Pollin, B., Rokyta, R. & Azerad, J. Reciprocal connections between the red nucleus and the trigeminal nuclei: a retrograde and anterograde tracing study. Physiol Res 47, 489–500 (1998).

46 Elbaz, M., Callado-Pérez, A., Demers, M., Kleinfeld, D. & Deschênes, M. A Vibrissal Pathway that Activates the Limbic System. under review (2021).

47 Zhu, H. & Roth, B. L. Silencing synapses with DREADDs. Neuron 82, 723–725, doi:10.1016/j.neuron.2014.05.002 (2014).

48 Gomez, J. L. et al. Chemogenetics revealed: DREADD occupancy and activation via converted clozapine. Science 357, 503–507, doi:10.1126/science.aan2475 (2017).

49 Mahler, S. V. & Aston-Jones, G. CNO Evil? Considerations for the Use of DREADDs in Behavioral Neuroscience. Neuropsychopharmacology 43, 934–936, doi:10.1038/npp.2017.299 (2018).

50 Manvich, D. F. et al. The DREADD agonist clozapine N-oxide (CNO) is reverse-metabolized to clozapine and produces clozapine-like interoceptive stimulus effects in rats and mice. Sci Rep 8, 3840, doi:10.1038/s41598-018-22116-z (2018).

51 Otchy, T. M. et al. Acute off-target effects of neural circuit manipulations. Nature 528, 358–363, doi:10.1038/nature16442 (2015).

52 Fetsch, C. R. et al. Focal optogenetic suppression in macaque area MT biases direction discrimination and decision confidence, but only transiently. Elife 7, doi:10.7554/eLife.36523 (2018).

53 Courville, J. The nucleus of the facial nerve; the relation between cellular groups and peripheral branches of the nerve. Brain Res 1, 338–354, doi:10.1016/0006-8993(66)90126-0 (1966).

54 Guest, J. M., Seetharama, M. M., Wendel, E. S., Strick, P. L. & Oberlaender, M. 3D reconstruction and standardization of the rat facial nucleus for precise mapping of vibrissal motor networks. Neuroscience 368, 171–186, doi:10.1016/j.neuroscience.2017.09.031 (2018).

55 Herfst, L. J. & Brecht, M. Whisker movements evoked by stimulation of single motor neurons in the facial nucleus of the rat. J Neurophysiol 99, 2821–2832, doi:10.1152/jn.01014.2007 (2008).

56 Kleinfeld, D., Moore, J. D., Wang, F. & Deschenes, M. The Brainstem Oscillator for Whisking and the Case for Breathing as the Master Clock for Orofacial Motor Actions. Cold Spring Harb Symp Quant Biol 79, 29–39, doi:10.1101/sqb.2014.79.024794 (2014).

57 Tsukahara, N. Sprouting of cortico-rubral synapses in red nucleus neurones after destruction of the nucleus interpositus of the cerebellum. (1974).

58 Murakami, F., Fujito, Y. & Tsukahara, N. Physiological properties of the newly formed cortico-rubral synapses of red nucleus neurons due to collateral sprouting. Brain Research 103, 147–151, doi:10.1016/0006-8993(76)90696-x (1976).

59 Alstermark, B. & Ekerot, C. F. The lateral reticular nucleus: a precerebellar centre providing the cerebellum with overview and integration of motor functions at systems level. A new hypothesis. J Physiol 591, 5453–5458, doi:10.1113/jphysiol.2013.256669 (2013).

60 Parenti, R., Cicirata, F., Panto, M. R. & Serapide, M. F. The projections of the lateral reticular nucleus to the deep cerebellar nuclei. An experimental analysis in the rat. Eur J Neurosci 8, 2157–2167, doi:10.1111/j.1460-9568.1996.tb00737.x (1996).

61 Qvist, H., Dietrichs, E. & Walberg, F. An ipsilateral projection from the red nucleus to the lateral reticular nucleus in the cat. Anat Embryol (Berl) 170, 327–330, doi:10.1007/BF00318738 (1984).

62 Freeman, J. H. & Steinmetz, A. B. Neural circuitry and plasticity mechanisms underlying delay eyeblink conditioning. Learn Mem 18, 666–677, doi:10.1101/lm.2023011 (2011).

63 McCormick, D. A. & Thompson, R. F. Cerebellum: essential involvement in the classically conditioned eyelid response. Science 223, 296–299, doi:10.1126/science.6701513 (1984).

64 Chapman, P. F., Steinmetz, J. E., Sears, L. L. & Thompson, R. F. Effects of lidocaine injection in the interpositus nucleus and red nucleus on conditioned behavioral and neuronal responses. Brain Research 537, 149–156, doi:10.1016/0006-8993(90)90351-b (1990).

65 Krupa, D. J., Weng, J. & Thompson, R. F. Inactivation of brainstem motor nuclei blocks expression but not acquisition of the rabbit’s classically conditioned eyeblink response. Behav Neurosci 110, 219–227, doi:10.1037//0735-7044.110.2.219 (1996).

66 Rosenfield, M. E. & Moore, J. W. Red nucleus lesions disrupt the classically conditioned nictitating membrane response in rabbits. Behav Brain Res 10, 393–398, doi:10.1016/0166-4328(83)90043-8 (1983).

67 Krupa, D. J., Thompson, J. K. & Thompson, R. F. Localization of a memory trace in the mammalian brain. Science 260, 989–991, doi:10.1126/science.8493536 (1993).

68 Giovannucci, A. et al. Cerebellar granule cells acquire a widespread predictive feedback signal during motor learning. Nat Neurosci 20, 727–734, doi:10.1038/nn.4531 (2017).

69 Heffley, W. & Hull, C. Classical conditioning drives learned reward prediction signals in climbing fibers across the lateral cerebellum. Elife 8, doi:10.7554/eLife.46764 (2019).

70 Larry, N., Yarkoni, M., Lixenberg, A. & Joshua, M. Cerebellar climbing fibers encode expected reward size. Elife 8, doi:10.7554/eLife.46870 (2019).

71 Kostadinov, D., Beau, M., Pozo, M. B. & Hausser, M. Predictive and reactive reward signals conveyed by climbing fiber inputs to cerebellar Purkinje cells. Nat Neurosci 22, 950–962, doi:10.1038/s41593-019-0381-8 (2019).

72 Moore, J. D., Deschenes, M. & Kleinfeld, D. Juxtacellular Monitoring and Localization of Single Neurons within Sub-cortical Brain Structures of Alert, Head-restrained Rats. J Vis Exp, doi:10.3791/51453 (2015).

73 Lavallee, P. et al. Feedforward inhibitory control of sensory information in higher-order thalamic nuclei. J Neurosci 25, 7489–7498, doi:10.1523/JNEUROSCI.2301-05.2005 (2005).

74 Kleinfeld, D. & Mitra, P. P. Spectral methods for functional brain imaging. Cold Spring Harb Protoc 2014, 248–262, doi:10.1101/pdb.top081075 (2014).

75 Bokil, H., Andrews, P., Kulkarni, J. E., Mehta, S. & Mitra, P. P. Chronux: a platform for analyzing neural signals. J Neurosci Methods 192, 146–151, doi:10.1016/j.jneumeth.2010.06.020 (2010).

76 Moore, J. D. et al. Hierarchy of orofacial rhythms revealed through whisking and breathing. Nature 497, 205–210, doi:10.1038/nature12076 (2013).

